# Exercise Myokine Irisin Enables Behavioral Stress Resistance in Mice

**DOI:** 10.64898/2026.07.14.737482

**Authors:** Simone M. Mellert, Joana F. da Rocha, Emily S. Levy, Juliet R. Freund, Jessica B. Fritz, Kaela Healy, Melanie J. Mittenbuhler, A Mu, Katherine A. Blackmore, Qiuyang Zhang, Michael V. Baratta, Christiane D. Wrann, Bruce M. Spiegelman, Benjamin N. Greenwood

**Affiliations:** Department of Integrative and Systems Biology, University of Colorado Denver, Denver, CO, 80217, USA; Cardiovascular Research Center, Massachusetts General Hospital and Harvard Medical School, Boston, MA, USA; Department of Psychology and Neuroscience, University of Colorado Boulder, Boulder, CO 80309, USA; Department of Psychology, University of Colorado Denver, Denver, CO, 80217, USA; Department of Cancer Biology, Dana-Farber Cancer Institute, Boston, MA 02115, USA; Department of Cell Biology, Harvard Medical School, Boston, MA 02115, USA; Department of Neurology, Mass General Brigham, Harvard Medical School, Boston, MA, USA; McCance Center for Brain Health, Massachusetts General Hospital, Boston, MA, USA

**Keywords:** social behavior, stress resilience, anxiety, BDNF, physical activity

## Abstract

Physical activity produces widespread benefits to physical and mental health, including protection against stress-related behavioral outcomes. However, the peripheral signals through which physical activity is translated into stress resistance remain incompletely understood. Irisin, an exercise-induced myokine cleaved from the transmembrane protein FNDC5 and released into circulation, has been implicated in improved cognition and neuroprotection, but its role in resistance to behavioral consequences of future adversity has not been examined. Here, we tested whether forced peripheral elevation of irisin is sufficient to protect against stress-induced behavioral outcomes. Adult male C57BL/6 mice received adeno-associated viral (AAV)-mediated peripheral expression of irisin or GFP control. AAV-irisin significantly increased circulating irisin levels, and six weeks later, mice were exposed to inescapable stress, a well-characterized model that produces anxiety-like behaviors, including reducing rodents’ natural inclination toward sociability. Elevated circulating irisin prevented the stress-induced reduction in social preference without altering general locomotor activity. Moreover, circulating irisin levels positively predicted individual differences in sociability. Additionally, peripheral irisin elevation increased brain-derived neurotrophic factor (Bdnf) expression, which also positively correlated with circulating irisin levels. Finally, we identified expression of the irisin receptor subunit integrin αV within the dorsal raphe nucleus, a key brain region implicated in the behavioral outcomes of inescapable stress that is modulated by prior exercise. Together, these findings identify peripheral irisin as a mediator sufficient to confer resistance to future stress, even in the absence of physical activity.

**Significance Statement:** Physical activity protects against the anxiety-like behavioral outcomes of future stress, but the peripheral mediators of this effect remain unclear. Here we show that forced peripheral elevation of circulating irisin, an exercise-induced myokine, is sufficient to produce behavioral stress resistance in mice. Irisin prevented the stress-induced reduction in social preference and positively correlated with individual differences in sociability. Forced expression of peripheral irisin also increased *Bdnf* expression in the brain, consistent with engagement of central neuroplasticity pathways that support adaptive behavioral responses. These findings demonstrate that an exercise-induced circulating factor can confer resistance to future stress, identifying irisin as a potential prophylactic target for stress-related mental health disorders.

## Introduction

Physical activity is a potent modulator of brain health, improving outcomes across a broad range of neuropsychiatric conditions, including age-related cognitive impairment (1–3), neurodegenerative disease (4,5), and mood disorders (6–9). These widespread effects highlight exercise as a behavioral intervention capable of improving mental health and reducing vulnerability to future pathology. Among these outcomes, aerobic exercise consistently reduces risk for development of future stress-related mental health disorders, such as anxiety and depression, by up to 25% (6,7,10). Given the widespread prevalence and recurrence of these conditions, identifying the mechanisms underlying exercise-induced stress resistance is critical. Elucidating these mechanisms could reveal novel prophylactic targets and inform therapeutic strategies for stress-susceptible individuals who are unable or unwilling to engage in regular physical activity.

Preclinical models are especially useful for studying the mechanisms underlying exercise-induced stress resistance because they allow precise control over individual history, including parameters of physical activity, stressor exposure, and behavioral outcome assessment. For example, rodents allowed voluntary access to running wheels are protected against the depression- and anxiety-like behavioral consequences of future stress (11). These types of preclinical studies reveal that exercise-induced stress resistance likely involves multiple central adaptations, including activity-dependent changes in gene expression, neurotrophic signaling, and neurotransmitter systems within stress-regulatory neural circuits (12). An emerging, yet understudied, area relevant to stress resistance involves peripheral signaling molecules that respond to physical activity and communicate with the brain.

Among exercise-induced signaling molecules, irisin has been identified as a key peripheral mediator of neural and behavioral adaptations. Irisin is released from skeletal muscle during physical activity through cleavage of the transmembrane protein FNDC5 in a *Pgc1α*-dependent manner (13). Exercise reliably increases circulating irisin across species, making the irisin pathway translationally relevant for studying exercise-induced adaptations. In mice, voluntary wheel running increases circulating levels of irisin (13) and in humans, mass-spectrometry studies confirm that plasma irisin is reliably detectable and increases significantly following aerobic exercise (14). Importantly, peripheral irisin crosses the blood-brain barrier (15,16) and is detectable in brain following peripheral elevation (26). Irisin binds αVβ5 integrin receptors, which have been identified as functional irisin receptors (18,19) and implicated in mediating irisin responses in brain tissue (20).

Accumulating evidence links irisin with exercise-induced cognitive improvements and neuroprotection (21–23). At the molecular level, exercise- and irisin-related benefits converge on brain-derived neurotrophic factor (*Bdnf*), a critical regulator of cognitive function and neural plasticity. Exercise rapidly increases hippocampal *Bdnf* (24), while exercise-induced *Pgc1α*/FNDC5 activation mediates hippocampal *Bdnf* expression (25). In mouse models of Alzheimer’s disease (AD), FNDC5/irisin is both necessary and sufficient to mediate the pro-cognitive effects of exercise on synaptic plasticity and memory (17). Importantly, forced peripheral expression of the mature, cleaved irisin via adenoviral vectors is sufficient, even in the absence of exercise, to increase hippocampal *Bdnf* expression, attenuate neuroinflammatory signaling, and improve cognitive performance in an AD mouse model (15). Peripheral elevation of irisin also confers protection against neurodegeneration, preserving dopaminergic neurons and improving motor outcomes in a mouse model of Parkinson’s disease (16) and mediates neuroprotection, including reducing neuronal loss, in a mouse model of multiple sclerosis (26). Together, these findings identify irisin as an exercise-induced peripheral signal that engages central plasticity-related mechanisms to influence both cognitive function and vulnerability to neurodegeneration.

In addition to the positive effects on cognitive function and brain health, irisin has also been implicated in mood regulation and emotional behaviors. In mice, subcutaneous recombinant irisin administered either intermittently over one month or acutely for five days reduces immobility in the tail suspension and forced swim tests and decreases anxiety-like behavior in the elevated plus maze, without affecting general locomotor activity in the open field (27,28); behavioral phenotypes identical to those reported following wheel running (29). Further, in rats exposed to chronic unpredictable stress, subcutaneous recombinant irisin mitigated stress-induced behaviors, reducing immobility in the forced swim test and restoring sucrose preference (30), in a manner similar to voluntary wheel running in the same stress model (31).

Although irisin influences emotional behaviors in both naïve rodents and those previously exposed to stress, reflecting a role in enhanced stress recovery, irisin has not yet been tested in a stress resistance model in which irisin elevation begins prior to stress. To address this gap, we examined whether increasing peripheral irisin could act prophylactically to prevent the behavioral consequences of future stress. We assessed stress-induced reductions in social preference as the primary behavioral readout, given that social avoidance is a hallmark across a broad range of stress-related psychiatric disorders (32–34).

Peripheral irisin was elevated using an adeno-associated viral (AAV8) vector expressing irisin, whereas AAV8-GFP served as a control. AAV8-irisin increased circulating irisin levels, and six weeks later mice were exposed to inescapable stress (IS), mirroring the duration of wheel running required to produce stress resistance. IS is a well characterized stress paradigm that reliably produces behavioral outcomes such as social avoidance (35,36). We hypothesized that IS would produce social avoidance in AAV8-GFP mice, but not in AAV8-Irisin mice, as a measure of behavioral stress resistance. Further, given the role of serotonin in the etiology and treatment of stress-related mental health disorders (37,38) and because exercise produces stress resistance by altering gene expression within serotonergic neurons of the dorsal raphe nucleus (DRN) and serotonin release in DRN targets (11,39–41), we examined whether the irisin receptor subunit αV is expressed in mouse serotonergic neurons of the DRN. We found that forced increases in peripheral irisin prevents IS-induced reductions in social preference, and that circulating irisin levels positively predict individual differences in sociability. Peripheral irisin also positively correlates with brain *Bdnf* levels, and we identified the αV subunit of the irisin receptor integrin αVβ5 in the DRN, providing anatomical evidence for a potential site through which irisin may modulate stress-related circuits.

## Results

### Peripheral irisin prevents stress-induced social avoidance without altering locomotion

To determine whether increasing peripheral irisin provides resistance to stress-induced social avoidance, mice received tail-vein injections of either AAV8-Irisin or AAV8-GFP control. Mice were exposed to IS or No Stress (NS) conditions 6 weeks later to mirror the timeframe required for males to attain stress resistance from exercise, followed by behavioral testing the next day (Fig 1A). AAV8-Irisin treatment increased plasma levels of irisin-FLAG, with no detectable levels in GFP controls (Fig 1B). Preference for social stimulus compared to nonsocial stimulus (social score) was significantly reduced by stress in GFP mice, but not in irisin mice. ANOVA revealed a significant main effect of virus (p = 0.05), but neither the main effect of stress, nor the virus × stress interaction were significant (Fig 1C). Because our analysis plan included a priori hypothesis testing, we performed targeted post hoc comparisons to confirm the predicted stress effect in GFP controls and to test the predefined hypothesis that irisin would protect against IS-induced social avoidance. As expected, IS reduced social interaction in GFP mice compared to NS (p = 0.03). In contrast, AAV8-Irisin prevented this effect: stressed mice given AAV8-Irisin had significantly higher social scores than GFP-IS (p = 0.01) and were comparable to both NS groups (p > 0.05; Figure 1C). These results indicate that AAV8-Irisin prevents the reduction in social preference produced by IS.

**Figure 1:**
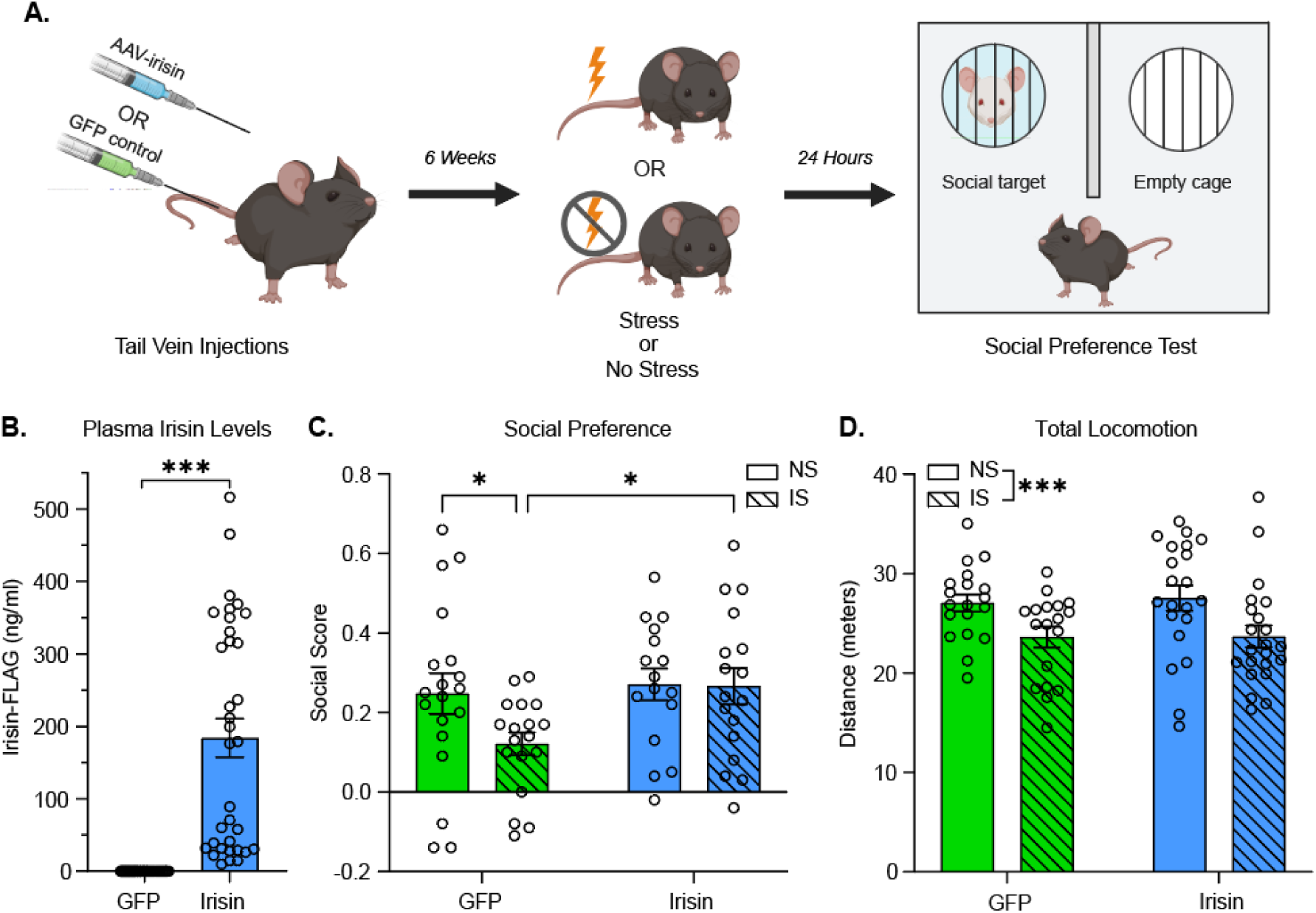
Effects of peripheral irisin and stress on social preference and locomotion. **A.** Experimental design created with BioRender.com. **B.** Irisin-FLAG plasma levels quantified via ELISA. Unpaired Welch’s t-test, ***pL<L0.001. **C.** Social score was significantly reduced by stress in GFP mice, but not in irisin mice. ANOVA, main effect of virus (F(1,66) = 3.89, p = 0.05), main effect of stress (F(1,66) = 2.37, p = 0.13), virus × stress interaction (F(1,66) = 2.04, p= 0.16). Social preference was also significantly reduced in stress GFP mice compared to stress irisin mice. Fisher LSD, *p < 0.05. **D**. ANY-maze tracking software calculated the total distance traveled for each social preference test. ANOVA, main effect of stress (F(1,76) = 10.91, p = 0.001), main effect of virus (F(1,76) = 0.06, p = 0.80), stress × virus interaction (F(1,76) = 0.04, p = 0.84). GFP-NS (N=19), GFP-IS (N=18), Irisin-NS (N=16), and Irisin-IS (N=17). For all, data are meanL±LSEM.

We next assessed total distance traveled to determine whether irisin altered general locomotor activity, as group differences in movement could influence social scores. Consistent with prior work (12), we found that stress significantly reduced the total distance traveled (p = 0.001; Fig.□1D), but distance traveled was not impacted by virus (Fig. 1D). Thus, AAV8-Irisin prevents stress-induced social avoidance despite a stress-induced reduction in locomotor activity.

### Circulating irisin levels correlate with sociability

We leveraged the heterogeneous range of circulating irisin produced by tail vein AAV injections to investigate a potential relationship between plasma irisin-FLAG levels and sociability. Linear regression analysis revealed that plasma irisin-FLAG levels positively predicted social score in all mice (p = 0.04; Fig 2A), but not total distance traveled (Fig 2B). Further automated measures of time spent in the social and nonsocial arenas were used to corroborate experimenter-scored social behavior. Consistent with the experimenter-scored social score in Fig. 1, plasma irisin-FLAG levels were significantly positively correlated with time spent in the social arena (p = 0.02; Fig 2C) and negatively correlated with time spent in the nonsocial arena (p = 0.002; Fig 2D). Collectively, these data show that peripheral irisin levels predict individual differences in sociability.

**Figure 2:**
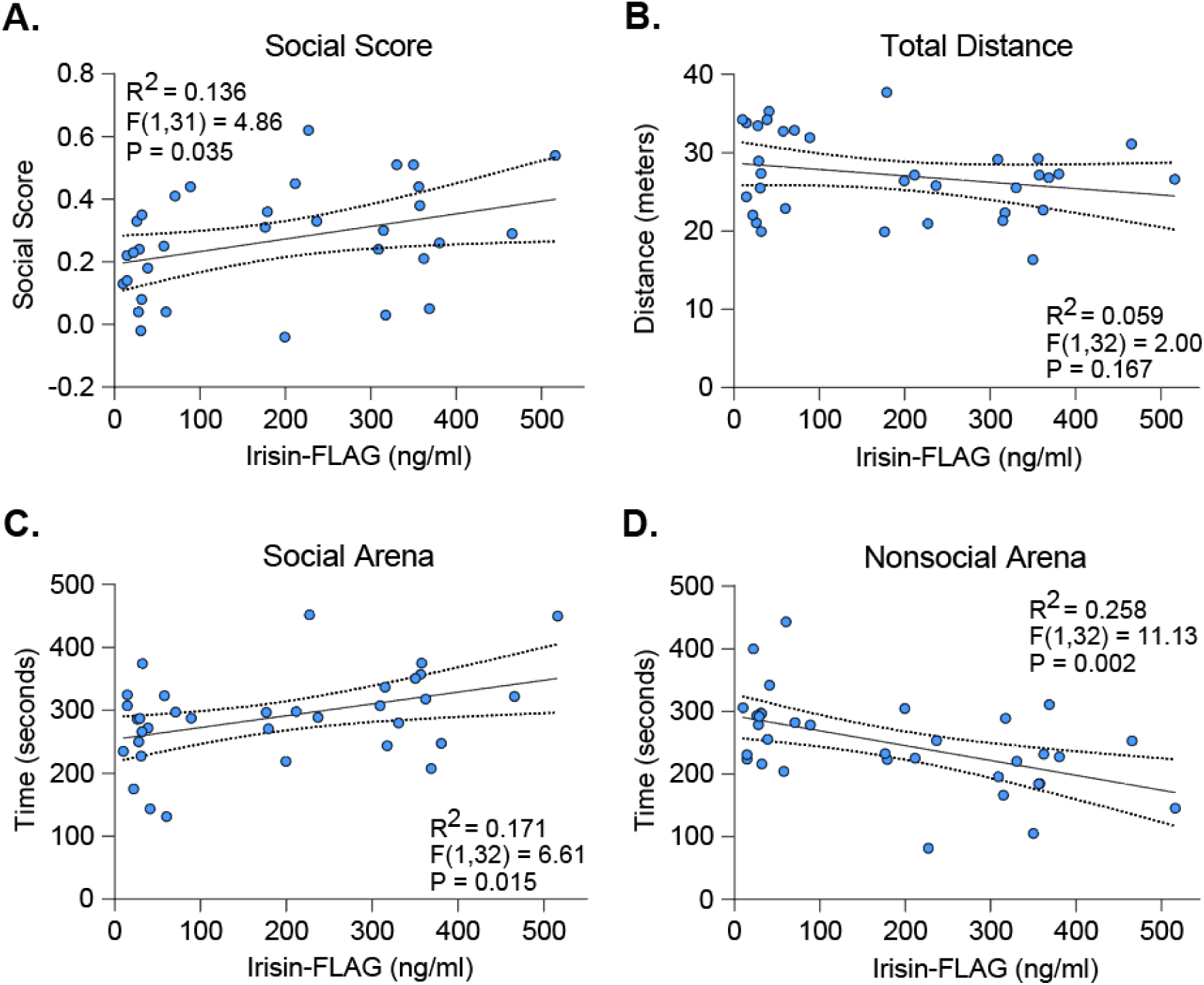
Irisin-FLAG plasma levels correlate with measures of sociability, but not locomotion. **A.** Irisin-FLAG plasma levels (ng/ml) positively correlate with social score. Linear regression analysis, (R^2^ = 0.14, F(1,31) = 4.86, p = 0.04). **B.** Irisin-FLAG plasma levels do not correlate with total distance traveled. Linear regression analysis, (R^2^ = 0.06, F(1,32) = 2.00, p = 0.17). **C.** Irisin-FLAG plasma levels positively correlate with time spent in the social arena. Linear regression analysis, (R^2^ = 0.17, F(1,32) = 6.61, p = 0.02). **D.** Irisin-FLAG plasma levels negatively correlate with time spent in the nonsocial arena. Linear regression analysis, (R^2^ = 0.26, F(1,32) = 11.13, p = 0.002). For all, N=33.

### AAV8-Irisin drives irisin expression in the liver without altering endogenous hepatic transcription

Prior work indicates that peripheral AAV8-Irisin predominantly targets the liver to increase viral irisin production (15). Liver tissue was collected for qPCR analysis to verify irisin expression and assess any broader hepatic transcriptional effects. As expected, AAV8-GFP treated mice had high levels of GFP mRNA in the liver whereas none was detected in AAV8-Irisin treated mice (Fig 3A), confirming effective expression of the control vector. Mice treated with AAV8-Irisin had high levels of irisin mRNA (Fig 3B). In contrast, endogenous *Fndc5* mRNA levels were unchanged between groups (Fig 3B), indicating that viral irisin did not alter native *Fndc5* expression.

**Figure 3:**
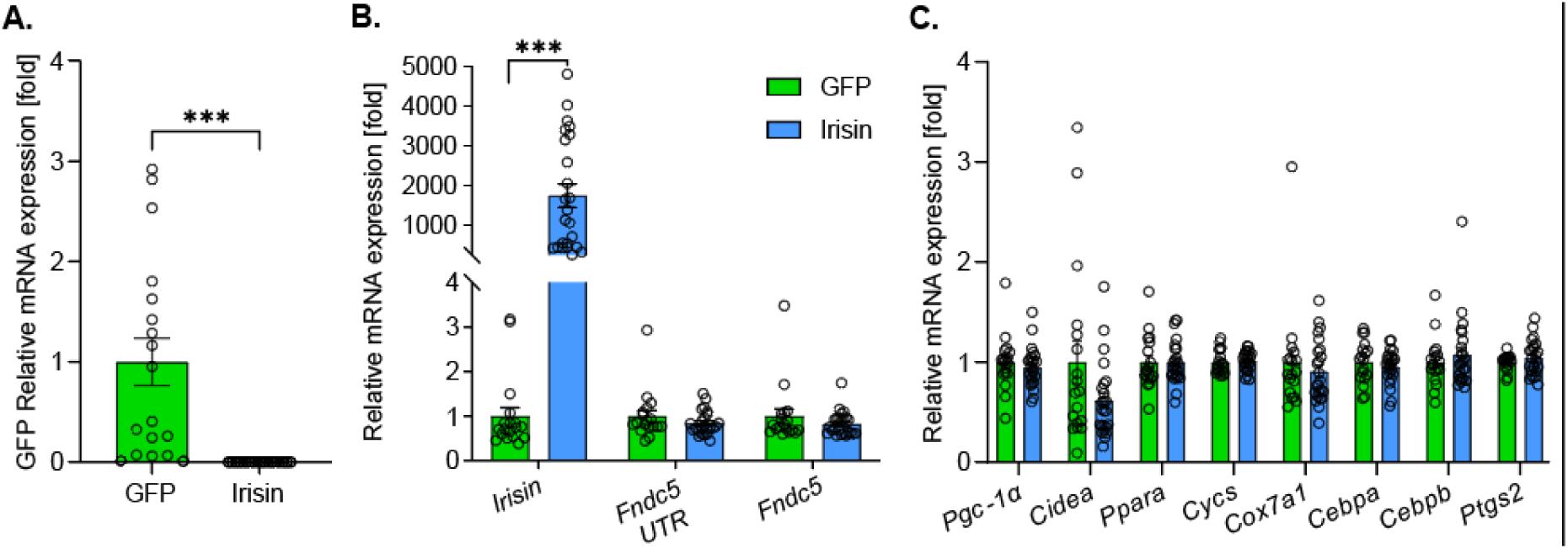
Peripheral irisin concentrates in hepatic tissue while maintaining normal transcript profile. **A.** qPCR analysis of hepatic GFP mRNA. unpaired Welch’s t test, p < 0.001. **B.** qPCR analysis of hepatic irisin and *Fndc5*mRNA. AAV8-Irisin mice had significantly higher *irisin* mRNA than GFP control mice unpaired Welch’s t test, p < 0.001, q < 0.001, while no difference was observed for *Fndc5* unpaired Welch’s t test, p = 0.27, q = 0.21 **C.** qPCR analysis of hepatic transcripts reveals no difference between irisin and GFP mice. Unpaired Welch’s t-test, p > 0.05, with FDR correction, q > 0.05 For all, GFP N = 18, irisin N = 24, and data are meanL±LSEM.

To evaluate whether forced peripheral irisin expression induces broader gene-expression effects in the liver, which could potentially indicate metabolic strain or toxicity, we quantified transcripts marking oxidative metabolism, mitochondrial function, and adipocyte differentiation (*Pgc1a, Cidea, Ppar*α*, Cytc, Cox7a1, Cebpa, Cebpb, Ptgs2*). Across all measured transcripts, no differences were detected between AAV8-Irisin and AAV8-GFP groups (Fig 3C), consistent with prior work showing that peripheral AAV8-Irisin delivery increases circulating irisin without impacting liver transcriptional profiles (15).This suggests that forced peripheral irisin expression is well tolerated in the liver and supports the translational feasibility of targeting peripheral irisin signaling for therapeutic applications.

### Peripheral irisin significantly increases Bdnf expression in the brain

To assess whether peripheral irisin influences endogenous central irisin transcripts, whole brain tissue was collected for qPCR analyses. Neither irisin mRNA nor endogenous *Fndc5* mRNA differed between groups (Fig 4A), suggesting that the AAV8-Irisin vector did not itself enter the brain, and that forced peripheral irisin expression did not alter endogenous central *Fndc5* expression.

**Figure 4:**
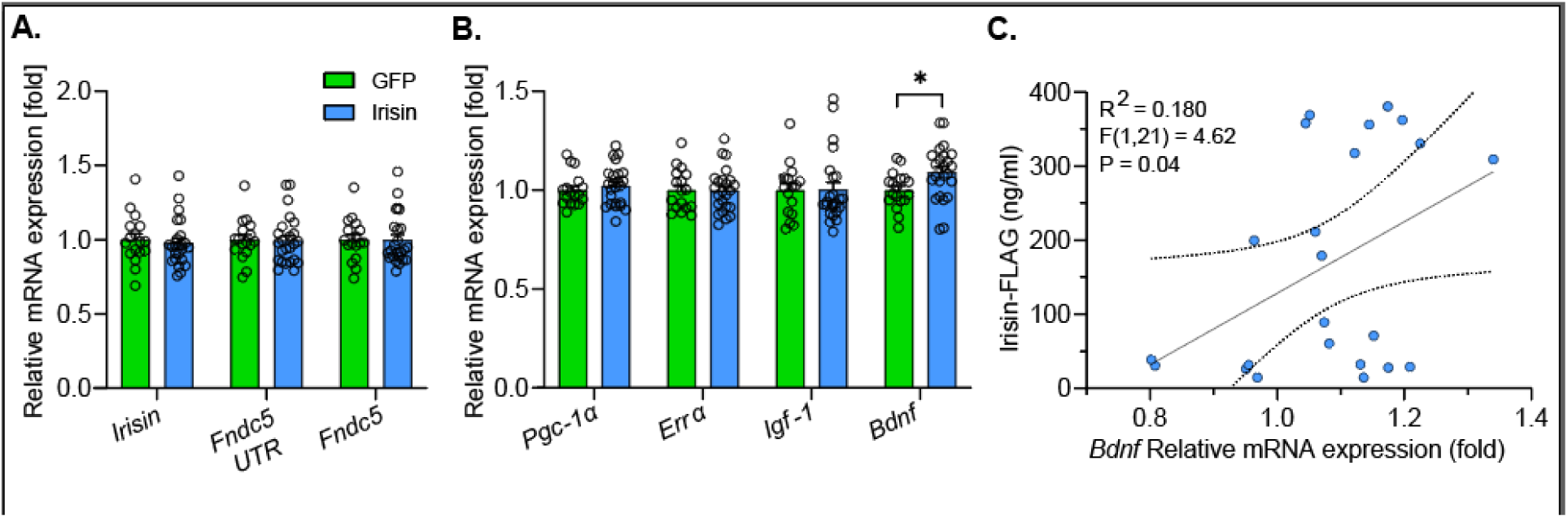
Peripheral irisin does not alter central *Fndc5* expression but does increase *Bdnf* mRNA expression. **A.** qPCR analysis of *Fndc5* mRNA in whole brain tissue reveals no effect of AAV8-Irisin relative to AAV8-GFP. Unpaired Welch’s t-test, p > 0.05, with FDR correction, q > 0.05.. qPCR analysis indicates significantly higher *Bdnf* mRNA expression in irisin mice compared to GFP mice; Unpaired Welch’s t-test, *p = 0.01, with FDR correction, q = 0.05, but not *Pgc-1*α, *Err*α, or *Igf-1*; unpaired Welch’s t test, p > 0.05, with FDR correction, q > 0.05. **C.** Plasma irisin-FLAG (ng/ml) positively correlates with central *Bdnf* mRNA expression. Linear regression analysis, R^2^ = 0.180, F(1,21) = 4.62, p = 0.04. For all, GFP N = 18, irisin N = 24, and data are meanL±LSEM.

We then examined whether peripheral irisin influences metabolic or neurotrophic markers in the brain. *Pgc*-*1*α and its transcriptional partner *Err*α were examined as upstream regulators of the FNDC5/irisin pathway in the brain, whereas *Bdnf* and *Igf-1* were assessed as exercise-responsive neurotrophic factors, with *Bdnf*, in particular, reported to be induced by FNDC5/irisin signaling in prior studies (22,25).

Expression of *Pgc-1*α, *Err*α, and *Igf-1* did not differ between AAV8-GFP and AAV8-Irisin treated mice (Fig. 4B). In contrast, *Bdnf* mRNA was significantly elevated in AAV8-Irisin mice compared to AAV8-GFP controls (Fig 4B), consistent with previous reports that peripheral irisin increases *Bdnf* expression in the hippocampus (25). To evaluate if this effect varied with circulating irisin levels, we conducted a linear regression analysis which revealed a significant positive correlation between plasma irisin and brain *Bdnf* expression (p = 0.04, Fig 4C). Overall, these data indicate that peripheral irisin engages plasticity-related gene expression in the brain, independent of engagement of endogenous central *Fndc5* or upstream regulatory pathways.

### Irisin receptor subunit integrin αV is present in a brain region critical for exercise-induced stress resistance

Since peripherally-induced irisin can be detected in the brain (15,16), we sought to determine if irisin has the potential to act directly within the DRN, a brain region critical for exercise-induced stress resistance (11,40). Using RNAscope, we assessed whether *Itgav* mRNA, encoding the αV subunit of the irisin receptor integrin αVβ5, is present in the DRN and whether it is co-expressed with *Tph2*, a marker of serotonergic neurons. *Itgav* mRNA was detected throughout the DRN at a density of 34.43 cells/mm^2^.

Within *Tph2* positive neurons (at a density of 278.24 cells/mm^2^), a small proportion of co-expression neurons (4.49 cells/mm^2^) were observed above background, as is consistent with detection of low-abundance transcripts at the single-molecule level (Fig 5). These data demonstrate that integrin αV is present within the DRN and is expressed in a small proportion of serotonergic neurons, supporting the anatomical capacity for irisin-mediated signaling in the DRN.

**Figure 5:**
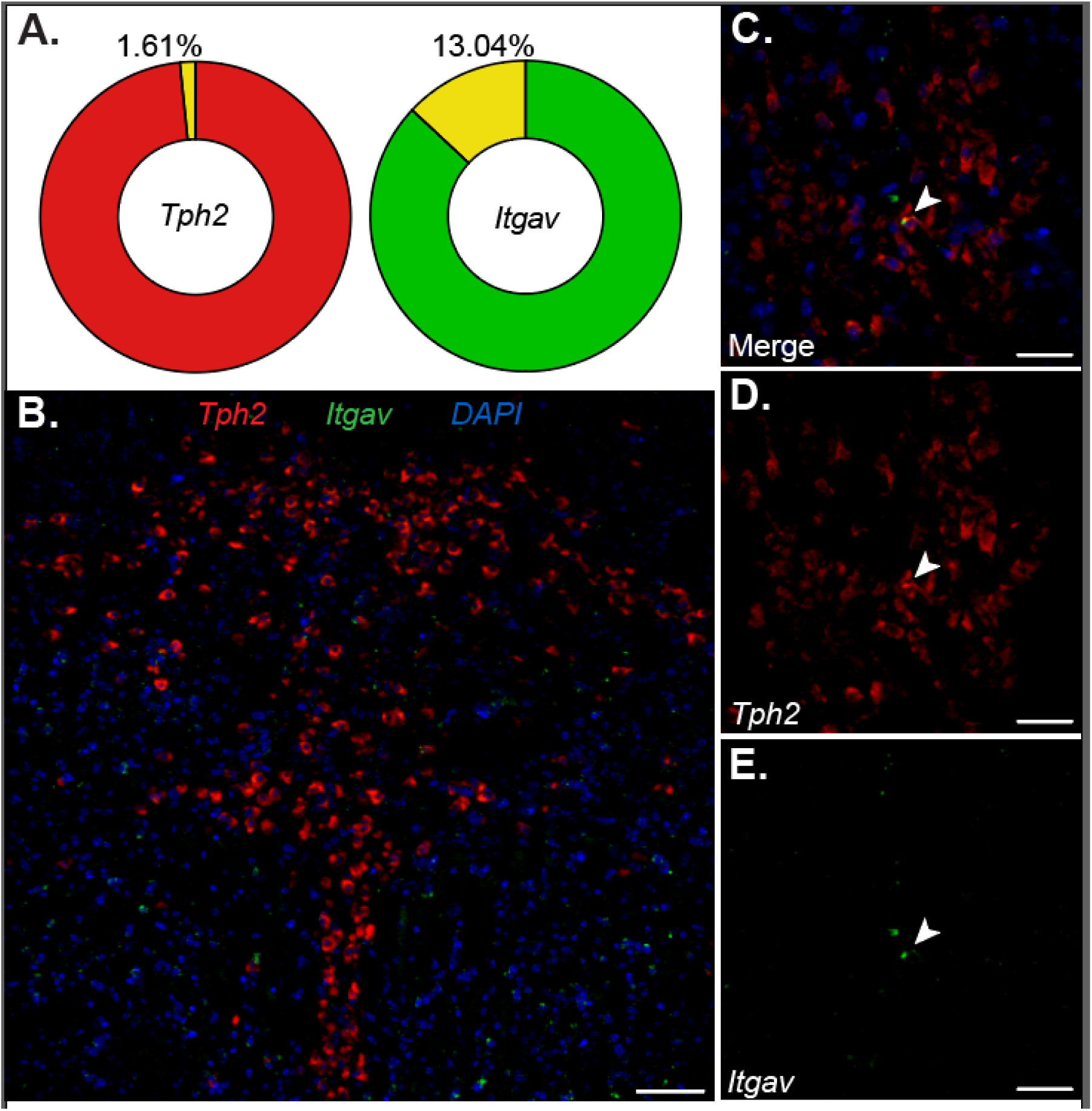
Integrin αV expression in the dorsal raphe nucleus (DRN). **A.** Quantification of percent tryptophan hydroxylase 2 mRNA (*Tph2,* 1.6%)- and Integrin αV (*Itgav,* 13.04%)- co-expression in the DRN. **B.** Fluorescent in situ hybridization labeling of *Tph2* (red) and *Itgav* (green) in DRN cells. Scale bar = 100 μm. **C.** Neuron co-expressing- **D.** *Tph2*- and **E.** *Itgav*- mRNA. Scale bars = 50 μ m.

## Discussion

The present study demonstrates that peripheral elevation of circulating irisin levels is sufficient to confer behavioral resistance to a future acute stressor, paralleling the established stress-protective effects of exercise. Using a validated model of IS-induced social avoidance (42,43), we show that mice with AAV8-mediated peripheral irisin expression are protected from future stress-induced reductions in sociability without alterations in general locomotor activity. Social avoidance is a core, transdiagnostic feature of stress-related psychiatric disorders that is associated with greater symptom severity and a lower likelihood of recovery from depression (33,34). In preclinical models, social behavior represents an ethologically relevant outcome, with stress-induced social avoidance reliably observed across multiple paradigms in mice (44,45). Because few interventions have been shown to prevent stress!Ilinduced social avoidance, these findings identify irisin as a unique peripheral signaling molecule with potential to promote proactive stress resistance against a core behavioral feature of stress!Ilrelated disorders.

Importantly, irisin expression was associated with higher social scores regardless of stress condition, and circulating irisin levels positively predicted individual differences in sociability. These data suggest that irisin may not simply preserve baseline behavior under adverse conditions but can actively bias behavior toward social approach. Automated scoring of time spent in the social and nonsocial arenas corroborated experimenter-scored social interaction, supporting the reliability of these findings. Further, the effects of irisin on sociability occurred independently of changes in general locomotor activity. Stress exposure significantly reduced locomotion, whereas peripheral irisin expression had no detectable effect, a pattern consistent with our prior findings in which stress reduces general activity regardless of exercise history (12). Together, these data suggest that forced peripheral irisin expression does not alter behavior by simply increasing activity or buffering against stress-induced hypoactivity, but instead selectively promotes social behavior.

The behavioral effects of peripheral irisin were accompanied by a significant increase in a central molecular marker associated with neural plasticity. AAV8-Irisin mice exhibited elevated whole-brain *Bdnf* expression relative to AAV8-GFP controls, consistent with prior work measuring hippocampal *Bdnf* (25). In addition, circulating irisin levels positively correlated with whole-brain *Bdnf*. Peripheral irisin elevation did not alter endogenous brain *Fndc5* expression, nor did it affect upstream transcriptional regulators *Pgc*-*1*α or *Err*α, indicating that peripheral irisin does not globally engage canonical *Fndc5*-dependent transcriptional pathways in the brain. Instead, the selective induction of *Bdnf* suggests that irisin preferentially engages activity-dependent neurotrophic signaling pathways.

Given that circulating irisin induced from AAV8-Irisin crosses the blood-brain-barrier (15–16, 26), the observed effects of irisin on brain and behavior observed here are most likely mediated by central effects. Indeed, emerging evidence suggests that peripheral irisin acts directly on neurons, inducing a neuroprotective transcriptional program and reducing neuronal loss in a mouse model of multiple sclerosis (26). These effects were accompanied by increased irisin binding and selective upregulation of αVβ5 integrin receptors in spinal cord neurons (26). Direct actions of irisin within stress-regulatory circuits would likely be mediated by receptor-based signaling pathways capable of influencing neuroplasticity-related gene expression. Consistent with this possibility, recent work showed that irisin-responsive signaling in hippocampal tissue depends on αVβ5 receptor activation, with receptor engagement activating focal adhesion kinase and downstream nitric oxide synthase pathways linked to increased *Bdnf* expression (47). This receptor-coupled signaling provides a rationale for examination of irisin receptor expression within stress-regulatory brain regions.

Stress-induced social avoidance is mediated by excessive activation of serotonergic neurons in the DRN during IS exposure (42). Compared to escapable stress, which fails to produce social avoidance, IS induces hyperactivation of DRN serotonergic neurons, resulting in increased local serotonin release that down-regulates 5-HT1A autoreceptors (48) and sensitizes serotonergic neurons to subsequent challenges such as social interaction (49). Exercise protects against stress-induced social avoidance by constraining DRN serotonergic activation during stress and preventing this sensitization (11,41). In this context, our identification of integrin□αV expression within the DRN provides a potential anatomical substrate through which irisin could influence stress-regulatory circuitry. Future studies are needed to determine how αV receptor signaling affects serotonin transmission during stress.

Although integrin αV is widely expressed throughout the brain, including the hippocampus, cortex, striatum, amygdala, and cerebellum (50,51), this is the first study to identify its expression within the DRN. While *Itgav* mRNA co-expression in *Tph2*-positive neurons was sparse, even low-abundance expression may be functionally significant. Moreover, most integrin αV signal was observed in non-serotonergic cells, raising the possibility that irisin acts through other DRN cell types, including astrocytes, which have been reported to express irisin receptors (52). A limitation of the current study is that αV can heterodimerize with several β subunits (β3, β5, β6, and β8). Because αVβ5 is the primary irisin receptor identified in peripheral tissues and hippocampal endothelial cells (18,47), future work should determine which αV-containing heterodimers are expressed in the DRN and whether β5 is co-expressed in serotonergic neurons. Despite this limitation, these results provide initial evidence that peripheral irisin may act within the DRN to influence stress-related circuitry.

To date, only a limited set of manipulations have been shown to confer resistance to IS, including behavioral control (53), voluntary wheel running (11,39), and prior ketamine administration (54). Among these, exercise is particularly translationally relevant as a freely accessible lifestyle factor whose stress-protective effects are conserved across species (55). Therefore, understanding how the beneficial effects of exercise are communicated from the periphery to the brain is especially important. Irisin has emerged as a key peripheral mediator of the beneficial effects of exercise. Irisin is reliably induced by aerobic exercise (13,14) and functions as a circulating signal capable of engaging central plasticity-related pathways in both rodents and humans (25,46). The present study demonstrates that peripheral elevation of irisin alone, in the absence of physical activity, is sufficient to prevent the reduction in social preference produced by future stress. Notably, circulating irisin levels not only prevented stress!Ilinduced reductions in social preference but were also positively correlated with increased sociability and elevated central *Bdnf* expression. These data extend prior work on irisin’s role in cognitive and mood-related behaviors to a proactive, stress-protective social domain. Overall, these findings provide evidence that an exercise-induced circulating factor can confer resistance to future stress, highlighting irisin as a potential prophylactic target for stress-related mental health disorders.

## Materials and Methods

### Animals and housing

Adult, male C57BL/6J mice (P42 on arrival; Jackson Laboratory, Bar Harbor, Maine) were housed in an AAALAC-accredited vivarium at the University of Colorado Boulder. Mice were group-housed (5 per cage), maintained at 22□°C with controlled humidity and a 12:12 h light/dark cycle (lights on at 07:00), with ad libitum access to food and water. Mice were allowed to acclimate to housing conditions for one week prior to experimentation and all testing occurred during the inactive (light) phase. All procedures were approved by the University of Colorado Boulder Institutional Animal Care and Use Committee and conformed to National Institutes of Health *Guide for the Care and Use of Laboratory Animals*. Only male mice were used to align findings with prior fundamental work on irisin (13,15).

#### Adeno-associated virus and tail vein injections

Production of the AAV8-Irisin-FLAG was performed according to Islam, 2021. Mice received intravenous injections of AAV8-GFP or AAV8-Irisin-FLAG (1□×□10^10^ - 2□×□10^10^ GC per mouse) into the tail vein under 1.5% isoflurane anesthesia. Virus was diluted in PBS to a final volume of 100□μl. Mice remained in their cages for 6 weeks after AAV8-GFP or AAV8-Irisin injections to match the duration of time necessary for exercise to enable protection against the behavioral consequences of IS in males (12,39).

#### Inescapable stress

Six weeks after injection of AAV8-GFP or AAV8-Irisin, mice were randomly assigned to a single session of IS or NS. IS mice were placed in a Plexiglas box (7 x 6 x 9 cm; Med Associates, St. Albans, VT) with a Plexiglas rod protruding from the rear of the box. The tail was secured to the rod with tape and affixed with copper electrodes and electrode gel. The session consisted of 100 tail shock trials (0.3 mA, 5 sec duration each) on a variable interval schedule (average ITI - 60 sec). IS mice were returned to their home cage immediately following the shock procedure. All mice were single-housed following IS or NS conditions. The IS procedure was used because prior work indicates that IS produces behaviors resembling symptoms of depression and anxiety (35,36,56), including social avoidance (43), and these behavioral consequences of IS are prevented by 6 weeks of prior voluntary wheel running (11,12).

#### Three-chamber social preference test

Twenty-four hours after IS, all mice were exposed to the three-chamber social preference test (43). Behavioral testing was conducted in a novel procedure room to minimize any shared cues between prior IS and behavioral testing environments. Mice were placed in the center compartment of an arena with three interconnected compartments (ANY-box, Stoelting, Wood Dale, IL). In one side, an empty sociability cage (Stoelting) was placed in the center, and the other side contained an identical sociability cage with a novel male conspecific, approximately 2 weeks younger than the experimental mouse. The experimental mouse was allowed to freely explore the entire arena for 10 min. Time spent investigating the social and nonsocial cages was later scored from videos by an experimenter blind to experimental conditions. Social investigation was defined as direct contact with the cage, including sniffing and rearing onto the apparatus. Social scores were quantified by the following formula: (social interaction time – nonsocial interaction time) / (social interaction time + nonsocial interaction time). Scores closer to 1 indicate a strong preference for the social stimulus and scores closer to 0 reflect reduced social preference. ANY-maze automated behavioral tracking software was used to quantify the total distance traveled and time spent in each arena. These are complementary measures such that time spent reflects arena preference or approach tendency while distance traveled helps distinguish social preference from changes in general activity and nonspecific arousal.

#### Perfusion and tissue collection

Four days after behavioral testing, mice were anesthetized under 1.5% isoflurane gas and cardiac blood was collected via the left ventricle into heparin-coated tubes. Samples were centrifuged to obtain the plasma fraction, which was aliquoted and stored at −80 °C until analysis. Whole brain and liver samples were snap-frozen in liquid nitrogen. Frozen brain tissue was subsequently cryo-pulverized and subdivided into homogeneous aliquots for downstream analysis. For RNAscope experiments, brains from a separate cohort of naïve adult male C57BL/6J mice (n = 9) were collected and fresh-frozen in isopentane cooled on dry ice between -30□°C to -40□°C prior to storage.

#### Irisin-FLAG ELISA

Irisin-FLAG levels in plasma samples and standards (0-200 ng/mL; AG-40B-0136-C010, Adipogen) were quantified by ELISA as described in Islam et al., 202 1(15). Briefly, 96-well plates were coated overnight, at 4 °C, with an anti-irisin capture antibody (MAB8880-100, R&D Systems) and blocked with 1% BSA for 1 hour at room temperature (RT). Following a 2-hour sample incubation, target binding was sequentially detected using an anti-FLAG antibody (14793S, Cell Signaling Technology) and a horseradish peroxidase (HRP)-conjugated secondary antibody (7074S, Cell Signaling Technology). Plates were washed between incubation steps. Colorimetric signal development was initiated with 3,3′,5,5′-tetramethylbenzidine (TMB) chromogen (ab171522, Abcam). Absorbance was measured at 450 nm using a FLUOstar Omega (BMG Labtech) plate reader after stopping the reaction with stop solution (ab171529, Abcam).

#### RNA isolation and qPCR

Total RNA was isolated from liver and brain using TRIzol reagent (Invitrogen) and purified using the RNeasy Mini Kit (Qiagen) according to manufacturer instructions, as described in Islam, 2021. First strand cDNA was synthesized using equal amounts of RNA and the High-Capacity cDNA Reverse Transcription Kit with RNase Inhibitor (Thermo Fisher, 4374967). Quantitative PCR was performed using Power SYBR Green PCR Master Mix (Thermo Fisher, 4367660) on a QuantStudio5 Real Time PCR System (Applied Biosystems). Relative quantification of gene expression normalized to Rps18 was determined by the comparative Ct method (ΔΔCT). Primer sequences are listed in Table 1.

**Table 1:**
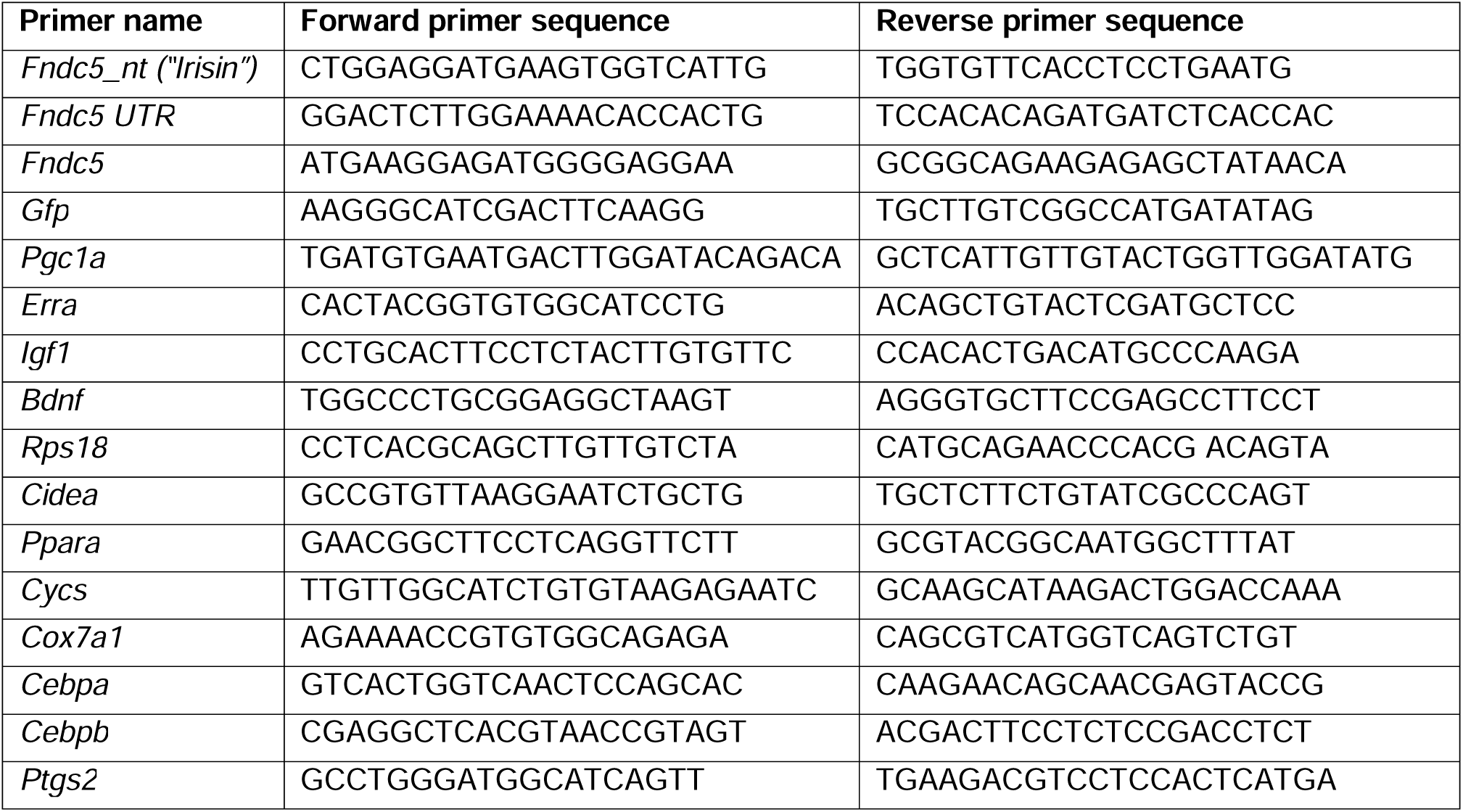
Primer Sequences.

#### RNAscope

Serial coronal sections (12□μm) containing the DRN were collected directly onto Superfrost Plus slides (Fisher Scientific) using a cryostat maintained at −24□°C. An RNAscope Multiplex Fluorescent Reagent v2 Kit was used according to the manufacturer’s instructions (Advanced Cell Diagnostics) for *Tph2* (Cat# 318691) and *Itgav* (Cat# 513901-C2) mRNA. *Tph2* was used to identify serotonergic neurons, as it produces the rate-limiting enzyme for central serotonin synthesis, and *Itgav* mRNA encodes the integrin αV subunit, which heterodimerizes with β5 to form the αVβ5 receptor. Opal dyes 520 and 570□nm (FP1487001KT, FP1488001KT; Akoya Biosciences) were matched to each probe, respectively. Slides were treated with DAPI (Advanced Cell Diagnostics) and coverslipped with antifade mountant (ProLong Diamond; Thermo).

#### Microscopy and quantification

Imaging double-fluorescent *in situ* hybridized tissue was acquired with a NikonA1 laser scanning confocal microscope and NIS-Elements software. The DRN region (three coronal sections for each subject, from Bregma (AP: -4.24 / -5.2mm) was imaged using a 10x/0.75NA objective with 405, 488, and 561□nm laser lines. Capture settings and z-stack image depth were kept constant across subjects. ImageJ/Fiji software was used to quantify the total number of cells expressing each label and, if any, their co-labeling. For each image, mean background fluorescence intensity was calculated as the average of four regions of interest placed within the medial longitudinal fasciculus, a DRN-adjacent white matter tract, and this value was subtracted from the image. Discrete pixels exceeding the defined background threshold were classified as positive signal. Co-labeled cells were identified as *Tph2* positive cells containing at least one *Itgav* mRNA punctum above background, consistent with RNAscope detection of low-abundance transcripts at the single-molecule level. We acknowledge the Light Microscopy Core Facility at the University of Colorado Boulder (RRID:SCR_018993).

#### Statistical Analyses

All statistical analyses were performed in GraphPad Prism (10.6.1). Following verification that data were normally distributed, ANOVA was used to compare group differences in social score, with virus (AAV8-GFP × AAV8-Irisin) and stress (NS × IS) as between-subject factors. Because our analysis plan included a priori hypotheses, planned post hoc comparisons (Fisher LSD) were conducted to confirm the expected IS-induced social avoidance in GFP controls and to test the predefined hypothesis that AAV8-Irisin would prevent this effect. Two social test statistical outliers were removed prior to analysis, yielding final sample sizes of GFP-NS (N=19), GFP-IS (N=18), Irisin-NS (N=16), and Irisin-IS (N=17). Total distance travelled was analyzed using ANOVA with the same between-subjects factors (virus and stress). Because no a priori comparisons were specified for locomotor activity, no planned post hoc tests were conducted for this analysis. Correlations were assessed using simple linear regression, with model fit summarized by R^2^, F statistic, and associated p values.

Pairwise group comparisons involving only two conditions (plasma irisin and hepatic GFP transcript expression) were analyzed using unpaired Welch’s t tests, which do not assume equal variances. qPCR analyses were performed in a subset of mice (GFP N = 18, irisin N = 24). For families of related transcript comparisons (liver and brain qPCR panels), unpaired Welch’s t-tests were used to obtain raw p values, which were then corrected for multiple comparisons using the Benjamini-Hochberg false discovery rate (FDR) procedure to yield adjusted q values. All tests were two-tailed, and effects were considered significant at p ≤ 0.05 and q ≤ 0.05 for FDR corrected transcript analyses.

To quantify *Itgav* mRNA expression within serotonergic neurons in the DRN, the percentage of *Tph2* positive cells co-expressing *Itgav* was compared across rostral, mid-rostral, and caudal subregions, as defined in Lowry, 2008 (57). ANOVA detected no differences among subregions, (F(2,11) = 0.23, p = 0.80); therefore, fluorescent positive cell counts (*Tph2*+, *Itgav*+, and co-labeled) were summed across the entire DRN (Bregma AP: -4.24/-5.2) within each mouse and then averaged across mice to obtain the total density of fluorescent positive cells per mm^2^. In all cases, data are reported as the mean ± SEM.

## Acknowledgments

This work was supported by NIH grants no. MH125898 (B.N.G.), NS087096 (C.D.W.), AG062904 (C.D.W.), the Alzheimer’s Association Research grant (C.D.W.), the Cure Alzheimer’s Fund (C.D.W.), and the Freedom Together Foundation (B.M.S.).

